# Combining metabolic engineering and fermentation optimization to achieve cost-effective oil production by *Cutaneotrichosporon oleaginosus*

**DOI:** 10.1101/2024.11.18.624102

**Authors:** Zeynep Efsun Duman-Özdamar, Rudolf Marcel Veloo, Elena Tsepani, Mattijs K. Julsing, Vitor A.P. Martins dos Santos, Jeroen Hugenholtz, Maria Suarez-Diez

## Abstract

Microbial oils, produced by oleaginous microorganisms, offer a sustainable alternative to plant-derived oils. Among these microorganisms, *Cutaneotrichosporon oleaginosus* attracts attention as a promising microbial cell factory for sustainable oil production due to its capacity to accumulate lipids with a similar composition to palm oil. Although *C. oleaginosus* can reach higher lipid contents than other oleaginous yeasts, suboptimal lipid yields and productivity limit economic feasibility. Enhanced productivity is necessary to have a feasible microbial oil production process via *C. oleaginosus*. In this study, we followed a combinatorial approach for strain design and bioprocess development to improve the lipid content and lipid yield. Initially, we deployed a full factorial design with genetic factors (ATP-citrate lyase (*ACL*), acetyl-CoA carboxylase (*ACC*), threonine synthase (*TS*)) and carbon-to-nitrogen ratio (C/N) in the medium. The C/N ratio appeared to have the most impact on oil accumulation. Combined with genetic modifications, lipid content and lipid yield increased by 1.6-fold. In a two-stage fermentation approach at a 2L scale, the triple transformant overexpressing *ACL*, *ACC*, and *TS* outperformed the wild-type by achieving a lipid content of 75.4% (w/w) with lipid productivity of 0.40 g L^-1^ h^-1^ and around 0.30 g lipids/g glycerol. In all, we established a cultivation strategy and strain that reached almost the theoretical maximum yield, and highest lipid content reported for a medium containing glycerol as a carbon source. These results strengthen the basis of using *C. oleaginous* as a platform for microbial oil production, thereby facilitating the development of processes substituting palm oil with a sustainable alternative.

**Highlights:**

- Full factorial design allowed selection of the best performer transformant.
- Overexpression of *ACL1, ACC,* and *TS* in *C. oleaginosus* provided higher lipid yield.
- Two-stage fermentation enhanced the lipid content of wild-type and transformant.

**Graphical Abstract:** 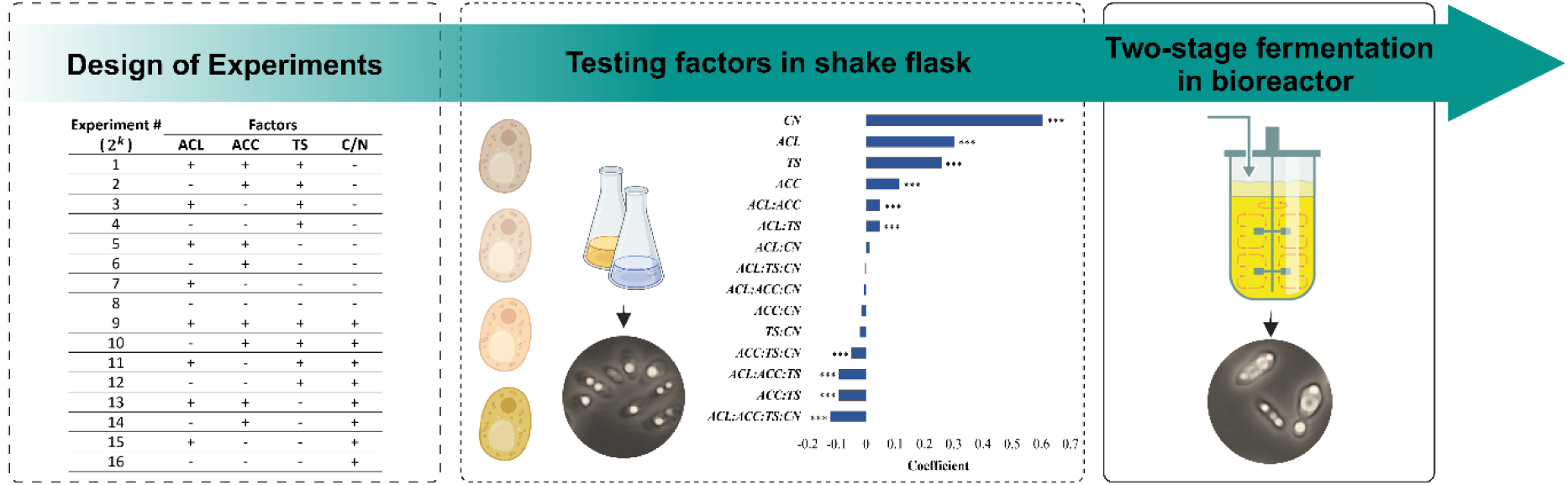

## 1. Introduction

Microbial oils, produced by oleaginous microorganisms, have emerged as a sustainable alternative to plant-derived oils and biodiesel production as they have a shorter life cycle, require less labor, show less affection by venue, season, and climate, and allow easy process scale-up [1,2]. Among the oil-producing microorganisms, oleaginous yeasts are particularly notable for their ability to accumulate more than 20% of their total biomass as lipids [3,4]. Furthermore, they are often considered superior for commercial applications due to their rapid growth, high lipid content, and high volumetric productivity [5]. These microorganisms store neutral lipids when there is excess carbon and other key nutrients, such as nitrogen, are limited [6]. Under these nutrient-limited conditions, cell growth is restricted, and oleaginous yeasts store excess carbon as lipids within intracellular lipid bodies [4,7].

Among these oleaginous yeasts, *Cutaneotrichosporon oleaginosus* (formerly known as *Apiotrichum curvatum, Cryptococcus curvatus, Trichosporon cutaneum, Trichosporon oleaginosus,* and *Cutaneotrichosporon curvatum*) stands out due to its high lipid accumulation capability, more than 40% and versatility in utilizing various carbon sources including glucose, xylose, cellobiose, sucrose, glycerol, lactose, and side streams such as crude glycerol and whey concentrate or permeate [8–12]. Under nitrogen limiting conditions, the produced fatty acid composition contains 25% palmitic acid (C16:0), 10% stearic acid (C18:0), 57% oleic acid (C18:1), and 7% 66 linoleic acids (C18:2), which is comparable to that of palm oil [13,14]. Additionally, it was presented as an effective fuel to replace both petroleum diesel and biodiesel produced from plant oils [15]. Due to these advantages, *C. oleaginosus* has been flagged as an attractive microbial-cell factory for industrial applications to sustain a bio-based circular economy. However, substantial optimization of the lipid production process is still needed to achieve high final microbial oil concentrations and lipid productivity for the implementation of microbial oil in industrial applications [16,17]. For *C. oleaginosus*, it has been calculated that the process is only economically viable if lipid accumulation of 60 % (w/w) and above is achieved, although it should be stressed that these calculations are based on glucose media and the use of well-controlled bioreactors [18].

Recently, research efforts have focused on designing *C. oleaginosus* as efficient cell factories by optimizing the medium and cultivation conditions and metabolic engineering to enhance lipid production [19–21]. The selection of different carbon and nitrogen sources and the C/N ratio appeared to influence the growth and lipid content of *C. oleaginosus* [22]. In our previous work, we reported that the lipid accumulation profile of *C. oleaginosus* was strongly dependent on the C/N ratio and temperature [14]. Moreover, we identified the optimum C/N ratio of the glycerol and urea-containing medium as 175 g/g. In addition to the medium composition, the selection of cultivation mode and feeding strategy have a significant impact on biomass, lipid yield, and process scalability. A two-stage fermentation approach, run in the fed-batch mode following the batch mode was reported as the optimal approach to achieve higher biomass densities and lipid yields thereby reducing the overall production cost [3,23]. In this approach, selecting the feeding strategy, such as feeding starting time points, and feeding modes, was reported as critical to avoid dilution and substrate inhibitions [3]. Meesters et al. (1996) studied two-stage fermentation and provided high cell densities of 118 g/L with 25% (w/w) lipid content and 0.11 g/g lipid yield in glycerol-containing medium [11]. Liang et al., (2010) utilized crude glycerol derived from yellow grease and achieved a biomass concentration of 32.9 g/l, with a 52% (w/w) lipid content [24]. Recently, we obtained improved lipid accumulation via the overexpression of ATP-citrate lyase (*ACL*), acetyl-CoA carboxylase (*ACC*), and threonine synthase (*TS*) [25]. ACL connecting the TCA cycle by converting citrate to acetyl-CoA and oxaloacetate, and ACC using acetyl-Co to produce malonyl-CoA are reported as key reactions to supply intermediates of fatty acid synthesis [26–28]. TS is responsible for threonine synthesis and is hypothesized to potentially increase the activity of the TCA cycle by pulling out the intermediate metabolites and balancing the overproduction of oxaloacetate [25,29].

In this work, we investigated the synergistic effect of these genetic factors (overexpression of *ACL, ACC, TS,* and a combination of these genes) and the C/N ratio of the medium by using a full-factor experimental design. The resulting regression model allowed us to capture the combinatorial effect of metabolic and environmental factors on lipid [30]. After evaluating the effect of genetic interventions and the C/N ratio, we selected the strain that resulted in the highest lipid yield and performed fermentation on a 2L scale using a two-stage fermentation approach in a controlled bioreactor. The wild-type and the best-performer transformant were tested for the lipid production, lipid yield, growth, and fatty acid composition of the produced microbial oil.

## 2. Materials and Methods

### 2.1. Strains, media, and growth conditions in shake-flask

*Cutaneotrichosporon oleaginosus* ATCC 20509, and transformants, ACL (ATP-citrate lyase overexpressed), and TS (threonine synthase overexpressed) [25], were used for genetic manipulation studies and maintained on Yeast extract Peptone Dextrose (YPD) agar plates containing 10 g/L yeast extract, 20 g/L peptone, 20 g/L glucose, 20 g/L agar. The maintained cultures were stored at 4 °C for up to a week. *Escherichia coli* Zymo 10B (Zymo Research, Orange, CA) was used for all cloning purposes throughout this study and maintained on Luria-Bertani (LB) agar (10 g/L tryptone, 10 g/L NaCl, 5 g/L yeast extract, 15 g/L agar) with ampicillin (100 μg/ml) or kanamycin (50 μg/ml) at 37°C.

The inoculum for shake flask experiments was prepared by transferring a single colony into 10 mL YPD broth (10 g/L yeast extract, 20 g/L peptone, 20 g/L glucose) in 50 mL tubes and incubated at 30 °C, 250 rpm for 18 h in a shaking incubator. Wild-type and other built *C. oleaginosus* transformants were cultivated in 300 mL shake flasks containing 60 mL minimal media consisting of glycerol as carbon source and urea as nitrogen source with set ratios of C/N 30 (g/g), and 175 (g/g) [14]. The C/N ratios were adjusted by changing the glycerol concentrations. Cultures were incubated at 30 °C, 250 rpm for 96 h in a shaking incubator. Cells were harvested at the end of incubation and centrifuged at 1780 g, 4 °C for 15 min. All shake flask experiments were performed in triplicates.

### 2.2. Plasmid assembly and preparation for transformation

Hygromycin B phosphotransferase gene (*HPT)* combined with the GPD promoter and GPD terminator, was cloned into a pUC57-Mini cloning vector as a backbone plasmid by GenScript Biotech (NJ, The US) (Table S1). pUC57-HPT was used as a selection marker for transformation studies. ACL and TS were used as background strains and pUC57-TS and pUC57-ACC were used to construct double transformants (Table 1) [25]. In order to build ACL_ACC_TS, ACL was used as a background strain. pUC57-TS-ACC plasmid was assembled. BsaI restriction sites were introduced to YATp-TS-YATt cassette by using YATp_BsaI_Fw and YATt_BsaI_Rv primers (Table S2) and assembly of the pUC57-TS-ACC was performed by using NEBridge Golden Gate Assembly Kit (BsaI-HF v2) (New England Biolabs, Ipswich, MA). The assembled plasmid was verified by whole plasmid sequencing (Eurofins, Germany).

**Table 1.** Plasmids and built strains.

| Strain | Background | Plasmids | Genotype | REF |
| --- | --- | --- | --- | --- |
| ACL | Wild-type | pUC57-ACL + pCG6-NAT | TEF1p-ACL-TEF1t + GPDp-NAT-GPDt | [25] |
| ACC | Wild-type | pUC57-ACC + pCG6-NAT | TPI1p-ACC-TPI1t + GPDp-NAT-GPDt |  |
| TS | Wild-type | pUC57-TS + pCG6-NAT | YATp-TS-YATt + GPDp-NAT-GPDt |  |
| ACL_TS | ACL | pUC57-TS + pCG6-HPT | TEF1p-ACL-TEF1t + GPDp-NAT-GPDt<br>YATp-TS-YATt + GPDp-HPT-GPDt | This study |
| ACL_ACC | ACL | pUC57-ACC + pCG6-NAT | TEF1p-ACL-TEF1t + GPDp-NAT-GPDt<br>TPI1p-ACC-TPI1t + GPDp-HPT-GPDt |  |
| TS_ACC | TS | pUC57-TS + pCG6-HPT | YATp-TS-YATt + GPDp-NAT-GPDt<br>TPI1p-ACC-TPI1t + GPDp-HPT-GPDt |  |
| ACL_TS_ACC | ACL | pUC57-TS-ACC + pCG6-HPT | TEF1p-ACL-TEF1t + GPDp-NAT-GPDt<br>YATp-TS-YATt-TPI1p-ACC-TPI1t + GPDp-HPT-GPDt |  |

Plasmids were transformed into *E. coli* Zymo 10B cells (Cat #T3020; Zymo Research, Irvine, CA, The US) by following the supplier’s instructions, and the transformed strains were stored at −80 °C as glycerol stocks. *E. coli* cells were grown overnight in 10 ml LB broth with 100 µg/ml ampicillin or 50 µg/ml kanamycin in 50 mL falcon tubes shaking at 250 rpm at 37 °C. Plasmid DNA was isolated using the GeneJET Plasmid Miniprep Kit (Cat #K0503; Thermo Fisher Scientific, MA, The US). Isolated plasmids were prepared for transformation by linearizing the pUC57-TS-ACC with HindIII and the other plasmids with EcoRI restriction enzyme according to the supplier’s instructions (NEB, Ipswich, MA, The US).

### 2.3. Co-transformation and selection for transformants

Electrocompetent cells were prepared as described earlier [25]. Co-transformation of linearized vectors containing the gene of interest and the plasmid containing the *HPT* gene was performed by adding approximately 1 µg per vector DNA into 50 µl electrocompetent cells and incubating them on ice for 5 minutes. Electroporation was performed using a pulse of 0.8 kVolt, 1000 Ohm, 25 µFarad (Bio-Rad, CA, The US) in 2 mm electroporation cuvettes. One mL of YPD broth was added immediately after pulsing. The cells were transferred to a 2 mL Eppendorf tube, and incubated for 2.5 hours at 30 °C, gently mixing the cells every 30 minutes by inversion. 100 µL cells were spread onto a YPD agar plate containing 100 µg/mL nourseothricin and 100 µg/mL hygromycin B for primary selection. The negative control was plated on YPD agar. Incubation of the plates was done at 30 °C for 48 hours. Grown colonies were randomly selected and streaked into a YPD agar plate including 100 µg/mL nourseothricin and 200 µg/mL hygromycin for secondary selection. Transformants were confirmed via colony PCR. The primers in Table S1 and DreamTaq Green PCR Master Mix (2X) were used by following the instructions from the supplier (Thermo Fisher Scientific, MA, The US, #K1081).

### 2.4. Experimental design for shake flask experiments and simulation

A full factorial design was employed by including ATP-citrate lyase (ACL), acetyl-CoA carboxylase (ACC), threonine synthase (TS), and C/N ratio as factors. This design resulted in 16 experiments (2^4^) performed in shake flasks (Table S4). Genetic factors were considered as categorical factors, the low level of genetic interventions was selected as ‘not modified’ and the high level was ‘overexpressed’ (Table 2). For the C/N ratio, C/N 30 was identified as low level and C/N175 was as high level.

**Table 2.**
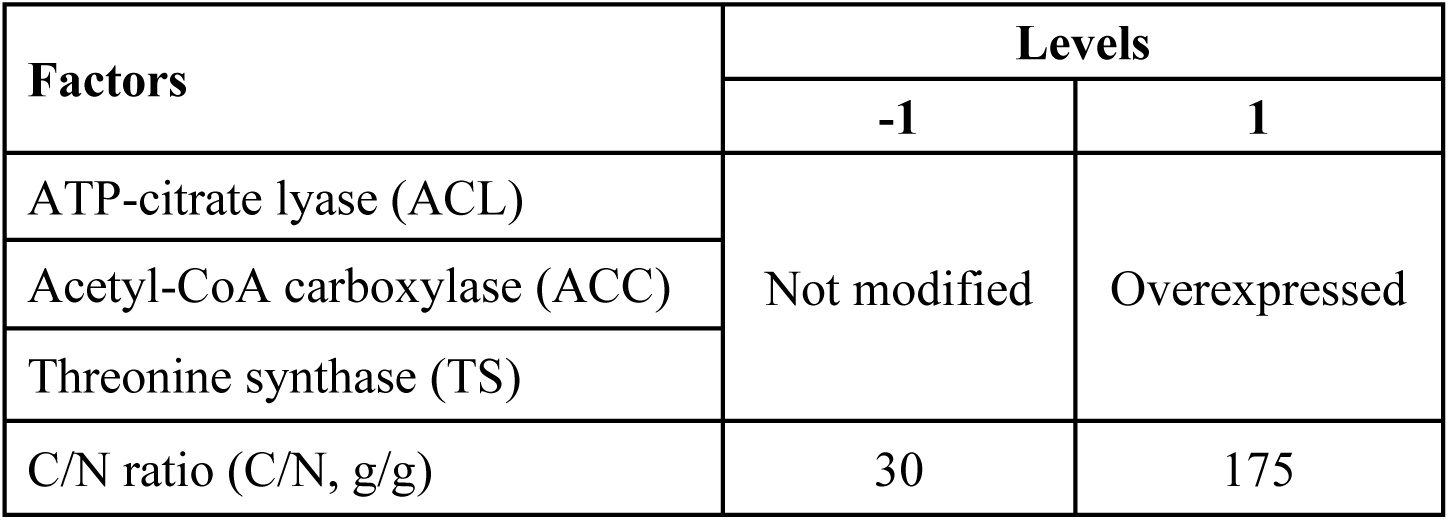
Genetic and medium composition as factors and their levels for the full factorial design.

All computational analyses were performed with R version 4.0.2 [31]. The relationship between the responses (*Y*), and factors (*x*) was expressed using the R lm function by fitting a linear model: *Y* = *β*_*o*_ + ∑ *β*_*i*_ *x*_*i*_ + ∑ *β*_*ij*_ *x*_*i*_ *x*_*j*_ *β_o_* represents the interception coefficient, *βi* is the linear coefficient (main effect), *β_ij_* is the interaction coefficient between factors *i* and *j.* The quality of the regression equations was assessed according to the coefficient of determination (R^2^). Statistical analysis of the model was performed using Analysis of Variance (ANOVA) and p < 0.05 was considered significant.

### 2.5. Bioreactor experiments

A 2L round-bottom bioreactor was used, equipped with a Rushton turbine, one marine impeller, and a control unit (Labfors 5, Infors HT, Switzerland). Inoculum was prepared by transferring a single colony into a 250 mL shake flask containing 50 mL YPG medium (10 g/L yeast extract, 20 g/L peptone, 16 g/L glycerol) and incubated at 30 °C, 250 rpm for 18 h in a shaking incubator. 18 ml overnight-grown culture of wild-type and ACL_ACC_TS was used as an inoculum for the 900 ml minimal medium in the 2L bioreactor. The medium contained 2g/L urea, 40g/L glycerol, 1g/L yeast extract, 2.7 g/L KH_2_PO_4_, 1.79 g/L NaH_2_PO_4_.7H_2_O, 0.2 g/L MgSO_4_.7H_2_O, 0.2 g/L MgSO_4_.7H_2_O, 0.1 g/L EDTA with the pH 5.5, 9 mL of 100X trace element solution (4 g/L CaCl_2_·2H_2_O, 0.55 g/L FeSO_4_.7H_2_O, 0.48 g/L citric acid.H_2_O, 0.1 g/L ZnSO_4_·7H_2_O, 76 mg/L MnSO_4_.H_2_O, 100 μL/L H_2_SO_4_ (36N)) and 200 µL of 100000X vitamin solution (2 mg/L biotin, 400 mg/L calcium pantothenate, 400 mg/L pyridoxine HCl, 400 mg/L thiamine HCl, 200 mg/L p-Aminobenzoic acid, 2 mg/L folic acid, 2 g/L inositol, 400 mg/L nicotinic acid, 100 mg/L riboflavin).

A two-stage fermentation approach was followed by starting in batch mode before switching to fed-batch mode. In the batch phase, the minimum dissolved oxygen (DO) concentration was set to 21% and the stirrer speed was set to a minimum of 300 rpm and automatically increased when DO was below the set point. The specific growth rates (µ, h^-1^) were determined by using the formula 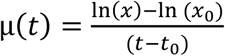 in which *x* is the dry cell weight at the end of the batch phase (for WT at 64 h and for ACL_ACC_TS at 48 h), *x_0_* is the initial dry cell weight. When glycerol was depleted, for which the increased DO level in the bioreactor was used as a signal, the fed-batch phase was initiated by feeding 70% (v/v) glycerol to the bioreactors. If sufficient glycerol was available in the medium, the DO level decreased rapidly due to the metabolic activities of cells. Increased DO levels activated the pump to provide a pulsed addition of glycerol. During the experiment, air was supplied at a rate of 0.5 l/min, the pH was kept constant at 5.5 by adding 10% (w/v) NaOH, and the temperature was kept at 30 °C. The pH, temperature, stirrer speed, and dissolved oxygen concentration were constantly monitored via Eve^®^ software (Infors HT, Switzerland). We collected 10 mL samples at 17h, 24h, 48h, 64h, and 72h, after that every 24h of the start of the fermentation. Bioreactor experiments were performed in duplicates.

### 2.6. Analytical methods

The growth of strains was monitored by measuring the OD_600_. Measured absorbance was converted into dry cell weight for shake flask experiments as explained by Duman-Özdamar et al. (2022) [14]. In bioreactor experiments, dry cell weight was measured by drying washed cells harvested on glass microfiber filter paper (Cytiva Whatman™, #1825-047, MA, The US) for 24 h in an oven at 105 °C.

The total fatty acids were determined quantitatively with a gas chromatograph (GC), 7830B GC systems (Aligent, Santa Clara, CA, The US) equipped with a Supelco Nukol^TM^ 25357 column (30m x 530 µm x 1.0 µm; Sigma-Aldrich, St. Louis, MO, The US), hydrogen as a carrier gas. Samples were prepared as described by Duman-Özdamar et al. (2022) [14]. Chloroform was evaporated under nitrogen gas and the remaining lipid in the tubes was dissolved in hexane before GC analysis.

Glycerol concentration was determined via HPLC analysis (Waters Alliance e2695, Milford, MA) with an RSpak KC811 column (ID = 8 mm, length = 300 mm, Shodex, NY) with a guard column RSpak KC-G (ID=6 mm and the length = 50 mm, Shodex, NY). The column was operated at 65°C with 3 mM H_2_SO_4_ as the mobile phase and a flow rate of 0.1 mL/min for 20 min. Peaks for components were detected and quantified with a refraction index detector (2414 RI Detector, Waters, Milford, MA). Peak integration and other chromatographic calculations were performed using Empower 3 software (Waters, Milford, MA). Identification and quantification of the corresponding compounds were achieved via comparisons to the standard curves (Figure S1). Urea concentration was determined semi-quantitatively by using Urea (BUN) Quick Test Strips (#MAS008, Sigma Aldrich, MO, USA).

## 3. Results

### 3.1 Selection of genetic and environmental factors

In our previous work, we observed the positive effect of overexpressing ATP-citrate lyase (*ACL*), acetyl-CoA carboxylase (*ACC*), and threonine synthase (*TS*) genes in *C. oleaginosus* on lipid content [25]. Additionally, various studies have reported the significance of the C/N ratio for obtaining improved lipid accumulation [14,22]. Herein we performed full factorial design to investigate the combinatorial effect of these genetic factors (ACL, ACC, TS) and the medium composition (C/N30: slightly N-limiting environment and C/N175: identified as optimum for wild-type) on total lipid, which resulted in total 16 experiments including 8 strains tested at selected C/N ratios in shake flasks (Table S3).

### 3.2 The impact of genetic and environmental factors on lipid production

Double (ACL_TS, ACL_ACC, TS_ACC), and triple (ACL_ACC_TS) transformants were built via co-transformation, and successful transformants were confirmed via colony PCR (Figure S2). After building the strains, the WT, single, double, and triple transformants were tested in C/N 30 and C/N 175 medium. At C/N30, ACL_TS accumulated 45.4% (w/w) lipids with 0.25 Y_P/S_ (g lipid/g glycerol), while WT provided 31.8 % (w/w) lipids with 0.18 g/g lipid yield (Figure 1A). Furthermore, ACL_ACC_TS outperformed all the other strains at C/N30 and provided a 1.6-fold increase in lipid content and lipid yield (51.8% (w/w), 0.29 g/g). This difference was observed also under the microscope between ACL_ACC_TS, which was more saturated with lipid bodies, and WT (Figure 1B). At a high C/N ratio, ACL_TS and ACL_ACC accumulated the utmost lipid content ∼60% (w/w), while ACL_ACC_TS accumulated 55% (w/w) lipids. ACL_ACC_TS strains showed the highest lipid yield of ∼0.29 (g lipid/g glycerol) at both media and improved the lipid yield by 1.6-fold compared to the WT. Moreover, while *TS* was overexpressed in single, double, or triple transformants, the obtained biomass concentration at C/N 30 increased by up to 34% and at C/N 175 by up to 22%. As a result of these increases, lipid weight (g/L) obtained via ACL_ACC_ TS transformant was boosted by 115% compared to WT.

**Figure 1.**
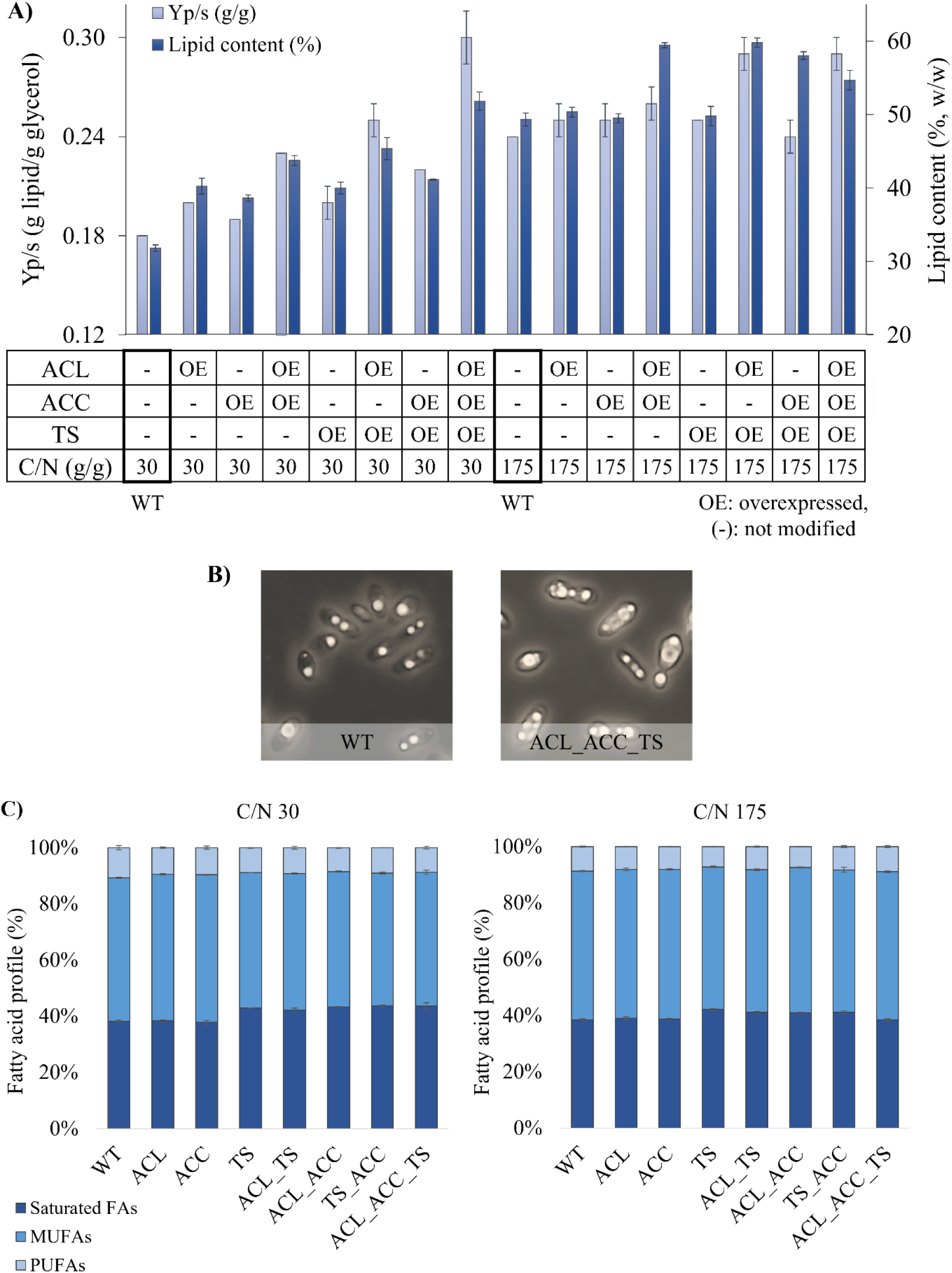
Effect of genetic targets and C/N ratio on lipid production and fatty acid profile. **A)** Lipid content and lipid yield on consumed glycerol (Y_P/S_, g/g) at different combinations of factors and levels. **B)** Microscope images of WT and ACL_ACC_TS growth at C/N 30 show that ACL_ACC_TS visibly became more saturated with lipid droplets (at 96h). **C.** Fatty acid profile (saturated fatty acids, PUFAs: polyunsaturated fatty acids, MUFAs: monounsaturated fatty acids) produced by WT and transformants at C/N30 and C/N175.

To further investigate the interactions between genetic factors and medium conditions and their main effects we fit a linear regression by focusing on the total lipid (g/L) as a response (Table S5). ANOVA was conducted to evaluate the statistical significance and suitability of the model and the quality of the model fit was assessed using the coefficient of determination (R^2^ = 99.34%), and the significance was confirmed via p-value: < 2.2e-16. The resulting equation highlighted that the C/N ratio of the medium showed the greatest impact followed by overexpression of *ACL, TS,* and *ACC* genes. Besides the main effects, we observed a significant positive interaction of ACL:ACC and ACL:TS on total lipid. On the other hand, there was no significant interaction between ACL:C/N, ACC:C/N, and TS:C/N.

In addition to the positive effect of these factors on lipid production, we observed that for both tested C/N ratios TS, double strains and triple strain showed higher content of saturated fatty acids with increased levels of C18:0 and C20:0 and the very long chain fatty acids (VLC-FAs) C22:0 and C24:0 (Figure 1C, Table S5). At C/N 30, saturated fatty acids varied between 38.2%-44%, while the content of monounsaturated fatty acids (MUFAs) and polyunsaturated fatty acids (PUFAs) was between 47.5%-52.6% and 8.5%-10.7%, respectively. When increasing the C/N to 175, we observed that the content of PUFAs shifted towards MUFAs for all strains. Moreover, at C/N175, ACL_TS, ACL_ACC, TS_ACC, and ACL_ACC_TS accumulated a higher content of unsaturated fatty acids with more MUFAs (C18:1) than at C/N30. Overall, the triple strain outperformed WT and the other transformants at high and low C/N ratios with the highest lipid yield, therefore, ACL_ACC_TS was selected for scale-up experiments.

### 3.3. Controlled conditions in bioreactor enabled higher lipid production

A two-stage fermentation approach was followed for ACL_ACC_TS, which outperformed the other strains, and WT in 2L bioreactors. Stage I was run in batch mode with the initial C/N around 15. When glycerol was depleted in the medium, stage II was initiated by pulsed feeding 70% (v/v) glycerol. The fed-batch phase was started for WT at 64 h and for ACL_ACC_TS at 48 h.

During the batch phase, the cell concentration of WT increased to 19.5 g /L (at 64h) and ACL_ACC_TS reached 21 g/L (at 48h) with a similar biomass yield on glycerol 0.53 g/g (Figure 2A). Although the biomass concentrations were comparable, ACL_ACC_TS represented a 1.35-fold higher specific growth rate (µ, 0.111 h^-1^) than WT (0.08 h^-1^). Through the end of the batch phase, the C/N ratio increased to around 90 g/g for both strains. While WT reached 5.3 g/L lipids with 27% (w/w) lipid content, we obtained 7.4 g/L lipids and 36.3% (w/w) lipid content with the triple strain at the end of the batch phase.

**Figure 2.**
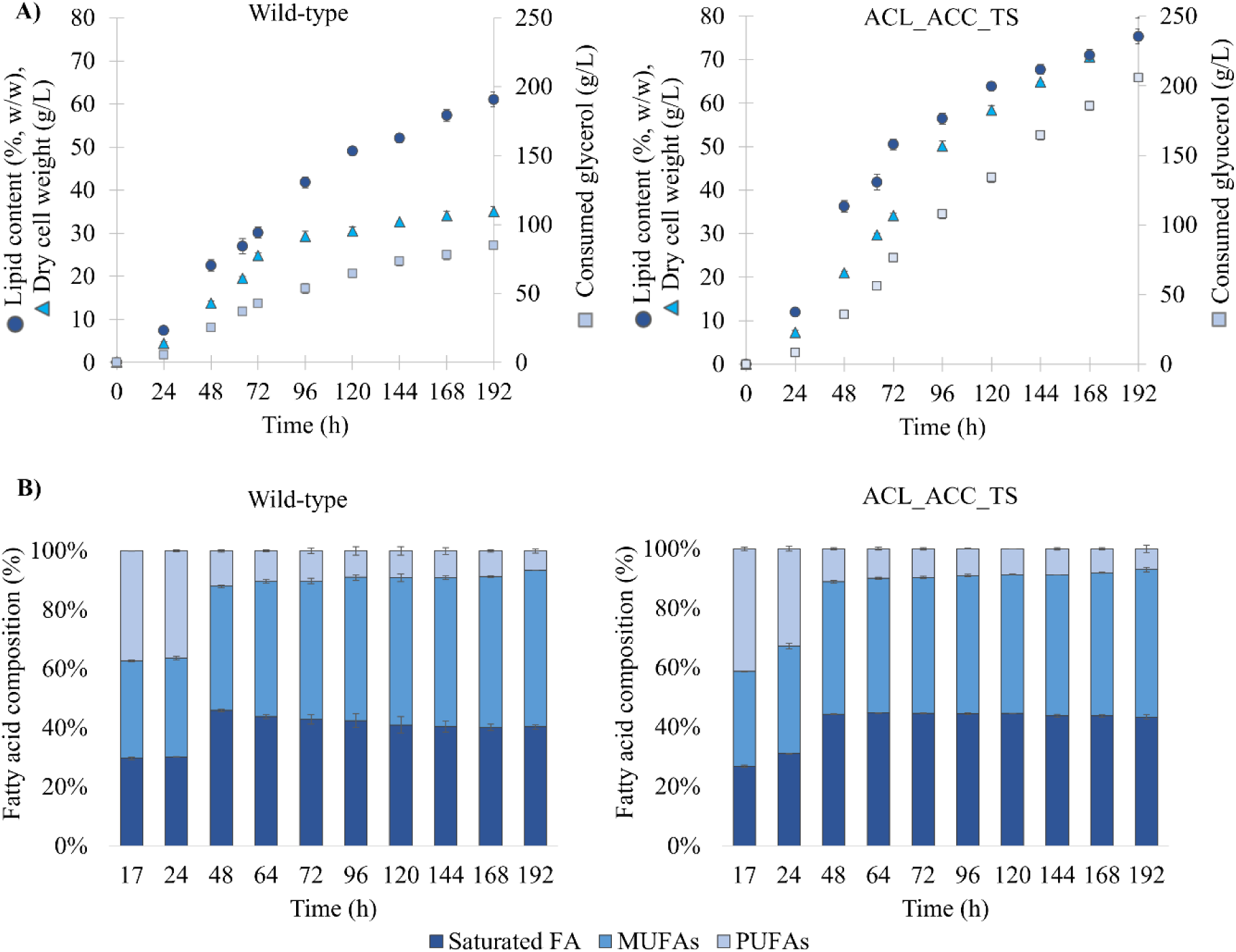
Characterization of ACL_ACC_TS and WT with 2L bioreactor experiments. **A)** Lipid content % (w/w) (dots), dry cell weight (g/L) (triangle), and consumed glycerol (square) during cultivation. **B)** Fatty acid composition (saturated fatty acids, PUFAs: polyunsaturated fatty acids, MUFAs: monounsaturated fatty acids) produced by WT and ACL_ACC_TS through two-stage fermentation. The batch phase was completed in 64 h for WT and 48 h for ACL_ACC_TS. The remaining fermentation process was then continued as a fed-batch until the end of 192 hours.

During the fed-batch phase, the biomass concentration of WT slowly increased to 35 g/L while the triple transformant reached a 2.2-fold higher biomass density (77.5 g/L) (Figure 2A). The lipid content of ACL_ACC_TS reached up to 75.4 % (w/w) with a yield of 0.30 g lipid /g glycerol and the lipid content of WT was 61% (w/w) with a yield of 0.25 g lipid/g glycerol. The lipid yield calculation in our work was performed based on consumed glycerol. In addition to glycerol, the cultivation medium employed in bioreactor experiments contained a yeast extract and vitamin solution that included a trace amount of carbon. However, this was excluded from calculations due to its negligible quantity in comparison to glycerol. The resulting total lipid produced by ACL_ACC_TS was 2.8-fold higher (61 g/L) compared to WT (22 g/L). Moreover, the average lipid productivity of triple strain was calculated as 0.40 g L^-1^h^-1^ and for WT as 0.18 g L^-1^h^-1^ at the fed-batch phase. Overall by using a two-phase fermentation approach, the lipid content of ACL_TS_ACC was increased by 38% and WT was improved by 25% compared to the shake flask experiments. This suggests that, in addition to the C/N ratio and genetic alterations, the development of a suitable process, such as a fed-batch approach can increase the overall yield of the production process.

Additionally, we investigated the fatty acid profile of WT and ACL_ACC_TS throughout the fermentation (Figure 2B). Before the start of the lipid accumulation phase (less than 20% lipids on biomass, until 24h), we observed that the major components were the PUFAs (C18:2 and C18:3). The content of C18:2 was stable and around 35% for WT in the first 24h. On the other hand, the content of C18:2 produced by triple strain decreased from 38% to 30% between 17h to 24h with accelerated lipid accumulation (Table S6-S7). From 48h of fermentation, we observed that the composition of PUFAs decreased to around 7% through the end of fermentation in both strains. MUFAs (C16:1, C18:1, C20:1, C22:1) increased from around 32% to 53.8% in WT and 50.3% in ACL_ACC_TS. Besides MUFAs, the content of saturated fatty acids (C16:0, C18:0) increased in WT from 30% to 41% and in ACL_ACC_TS from 27% to 44%. Thus ACL_ACC_TS produced higher contents of saturated fatty acids that were also observed in the shake flask experiments. The switch from the batch phase to the fed-batch phase did not lead to significant changes in the fatty acid composition of the WT and ACL_ACC_TS. Overall, the selected genetically modified transformant, ACL_ACC_TS, outperformed the WT also in a controlled bioreactor set up at a technical scale representative for larger-scale microbial oil production.

## 4. Discussion

In this study, we investigated the potential of *C. oleaginosus* for producing microbial oils on a glycerol-containing medium by combining strain engineering and a bioprocess development approach. The critical role of ACL and ACC reactions in providing precursor molecules for fatty acid formation and the positive effect of TS overexpression on fatty acid synthesis was reported for *C. oleaginosus* [25,28]. We showed, by applying a full factorial design, that the overexpression of ACL, ACC, TS, and high C/N ratio significantly enhanced the lipid content at the shake flask level. The regression analysis confirmed the significant impact of the C/N ratio on lipid production, followed by the overexpression of ACL, TS, and ACC genes. Although, regression analysis reflected a negative impact of ACL:ACC:TS interaction, the coefficient is much less compared to the main effects. Additionally, we obtained the highest lipid yield with ACL_ACC_TS at both C/N ratios, and we observed a significant positive interaction between ACL:ACC, as well as ACL:TS. Possibly, the combination of these genetic modifications synergizes to maximize the flow of carbon toward lipid biosynthesis.

After confirming the potential of ACL_ACC_TS at low and high C/N ratio medium we performed bioreactor experiments with WT and triple strain by following a two-stage fermentation strategy. The ACL_ACC_TS strain showed superior performance with a lipid content of 75.4% (w/w) and average lipid productivity of 0.40 g L^-1^h^-1^, compared to the WT (61% w/w and 0.18 g L^-1^h^-1^). Meesters et al. (1996) studied lipid accumulation of *C. oleaginosus* in a glycerol-containing medium using a fed-batch mode and obtained 25% (w/w) lipid accumulation with a yield of 0.11 g lipid/g glycerol [11]. To our knowledge, we obtained the highest lipid content for *C. oleaginosus* on a glycerol-containing medium. Furthermore, triple strain resulted in ∼0.30 g lipids/g glycerol, which is the highest lipid yield reported for *C. oleaginosus* grown in glycerol, and almost equal to the reported theoretical maximum yield on glycerol [4]. In addition to lipid productivity, ACL_ACC_TS grew with a higher specific growth rate in the batch phase and reached higher biomass concentrations at the end of the fed-batch phase compared to WT. The enhanced growth profile of the triple transformant may be linked to the overexpression of the TS gene, which potentially increases the intracellular amino acid availability. An elevated level of threonine synthesis has been demonstrated to enhance the growth rate of *Escherichia coli* [32] the alleviation of uracil and leucine auxotrophy has been shown to improve the growth rate of *Yarrowia lipolytica* [33].

The fatty acid composition of the lipids produced by ACL_ACC_TS contained a slightly higher content of saturated fatty acids compared to WT. In the first 24 hours, batch mode, cells were at the growth phase and accumulated higher content of C18:2 and C18:3 for both strains [11]. During the fed-batch phase, the fatty acids composition represented a similar pattern. This composition is comparable to the previously reported fatty acid compositions produced by wild-type [11,34] and also to the composition of palm oil thus representing a great alternative to plant-based oils for biodiesel production [13,15].

Altogether, we established a cultivation strategy on a glycerol-based medium and a strain that can reach above 75% (w/w) lipid content and higher biomass concentration resulting in 61 g lipid/L compared to WT. These results emphasize the potential of *C. oleaginosus* also for the bio-valorization of crude glycerol into single-cell oil [35–40]. Integrating biodiesel-derived glycerol for yeast oil production will further reduce cultivation costs and sustain a circular bio-economy by reusing the glycerol side stream thereby contributing to the development of processes substituting plant-derived oils with a sustainable alternative.

## Conclusion

In this study, we appraised the combinatorial effect of ACL, ACC, TS overexpression, and C/N ratio as factors in the lipid production of *C. oleaginous* via implementing the full factorial design. The ACL_ACC_TS transformant outperformed others, improving lipid content by 1.6-fold and reaching the highest lipid yield in shake flask experiments. Following shake flask experiments, we characterized ACL_ACC_TS, and WT at 2L bioreactors by following a two-phase fermentation approach. DO-coupled glycerol feeding following a batch stage improved the lipid content of WT to 61% (w/w) and of ACL_ACC_TS to 75.4% (w/w). The ACL_ACC_TS strain showed superior performance, achieving a 2.8-fold higher lipid production, an average lipid productivity of 0.40 g L^-1^ h^-1^, and a lipid yield of around 0.30 g/g surpassing the WT. This represents the highest lipid yield reported for *C. oleaginosus* on a glycerol-containing medium, approaching the theoretical maximum yield on glycerol. Our study underscores the potential of integrating genetic modifications with bioprocess optimization to significantly enhance lipid production in *C. oleaginosus*. These findings contribute to the development of economically viable microbial oil production processes, supporting the transition towards a sustainable bio-based circular economy.

## Author contributions

All authors conceived and designed the study. ZEDÖ and MSD performed the data analysis. ZEDÖ, ET, and RMV performed the experiments. ZEDÖ drafted the manuscript. VAPMdS, JH, and MSD acquired project funding, and VAPMdS, JH, MSD, and MKJ conceived and supervised the research. All authors reviewed and edited the study. All authors read and approved the final manuscript.

## Acknowledgments

The graphical abstract was created with BioRender.com.

## Supplementary Materials

**Table S1.**
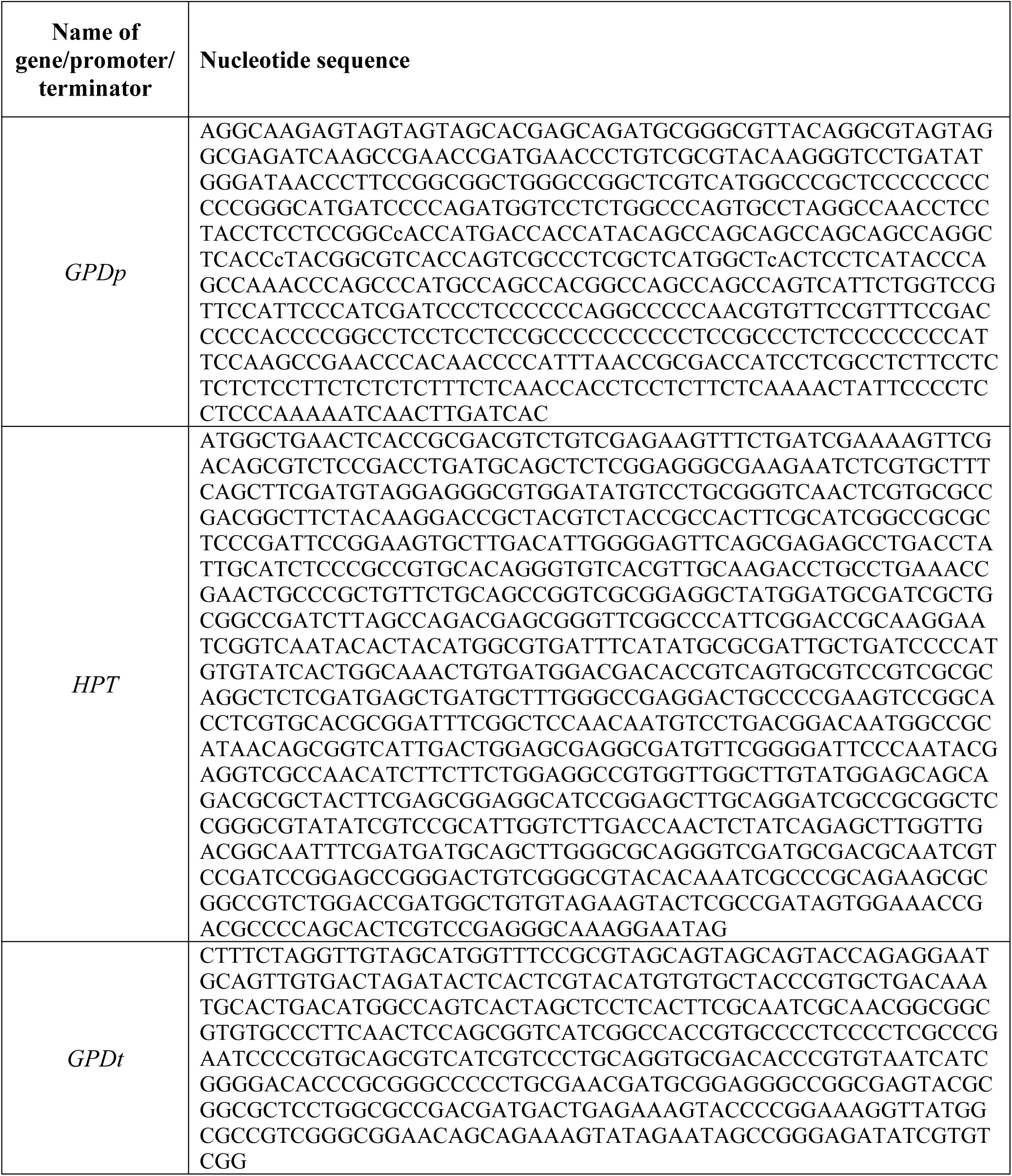
Nucleotide sequences of promoters, genes, and terminators.

**Table S2.**
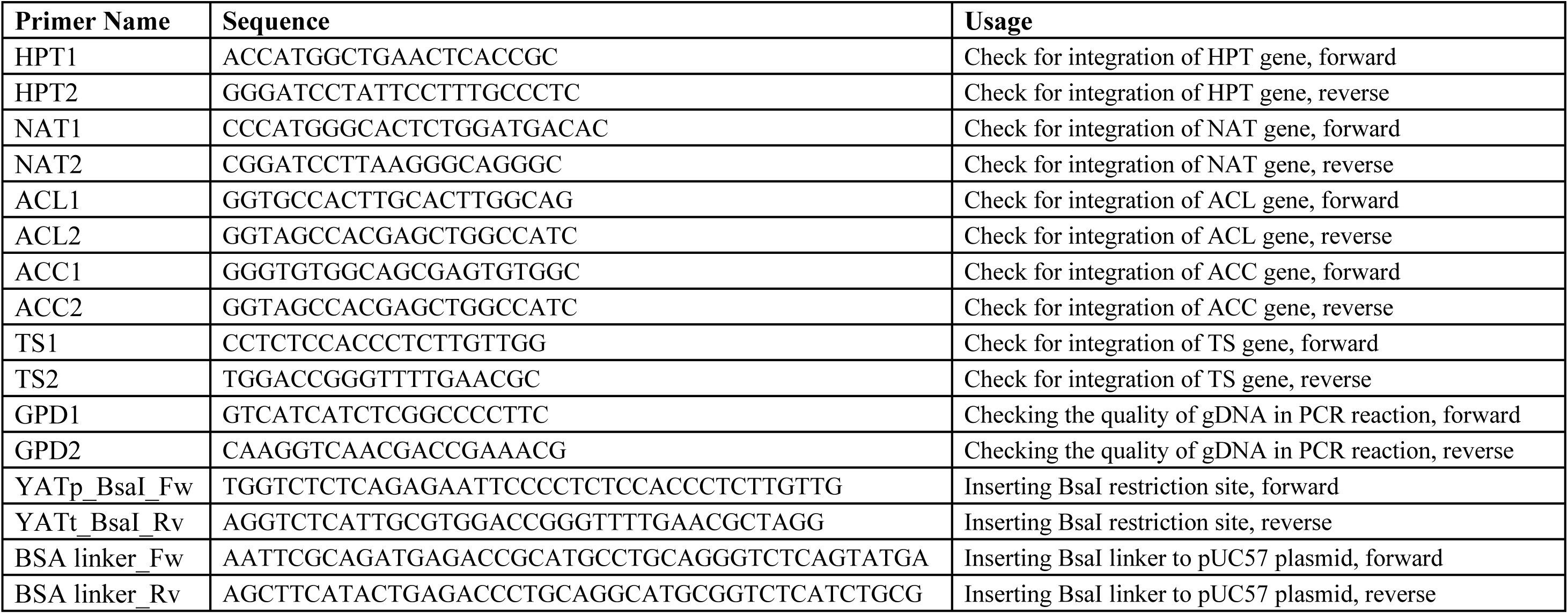
Primers designed for colony PCR and insertion of BsaI restriction sites.

**Figure S1.**
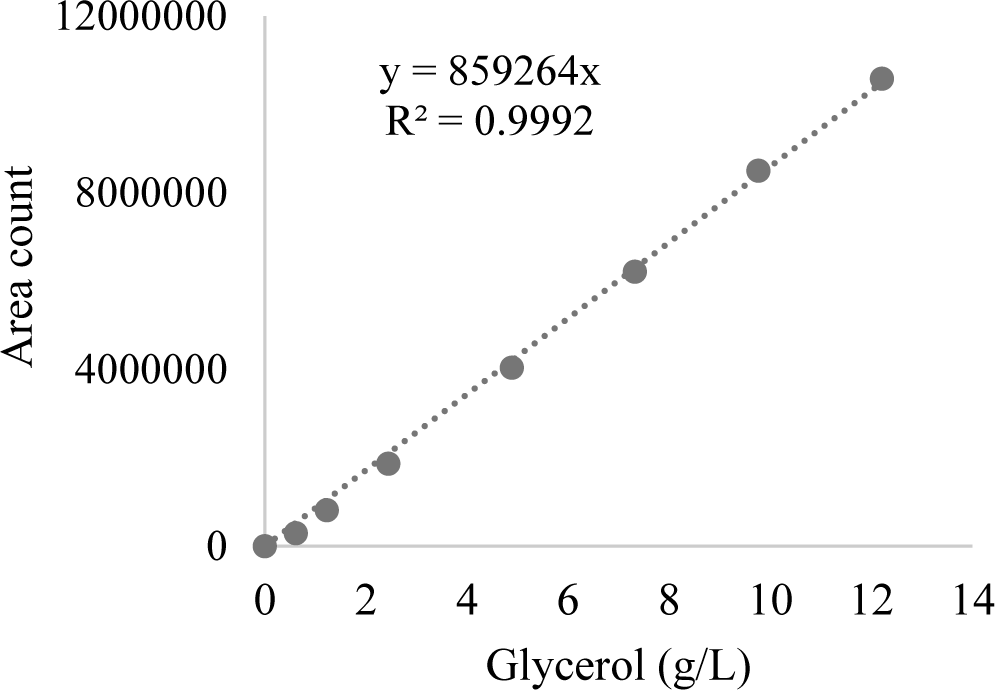
Calibration curve of glycerol for calculating the glycerol concentration of medium.

**Table S3.** Experimental runs and results of full factorial design. Dry cell weight, lipid content, and lipid yield on glycerol (Y_P/S_, g lipid/g consumed glycerol) of WT, at C/N 30 and 175 minimal medium (at 96h).

| Run | Factors (coded values) |  |  |  | Strain | C/N<br>(g/g) | Dry cell<br>weight<br>(g/L) | Lipid content<br>(%, w/w) | Y<sub>P/S</sub><br>(g lipid/g<br>consumed<br>glycerol) |
| --- | --- | --- | --- | --- | --- | --- | --- | --- | --- |
|  | ACL | ACC | TS | C/N |  |  |  |  |  |
| 1 | -1 | -1 | -1 | -1 | WT | 30 | 4.16 ± 0.23 | 31.81 ± 0.46 | 0.18 ± 0 |
| 2 | 1 | -1 | -1 | -1 | ACL |  | 4.57 ± 0.08 | 40.26 ± 1.08 ** | 0.2 ± 0 |
| 3 | -1 | 1 | -1 | -1 | ACC |  | 4.41 ± 0.16 | 38.63 ± 0.45 ** | 0.19 ± 0 * |
| 4 | 1 | 1 | -1 | -1 | ACL_ACC |  | 5.12 ± 0.08 ** | 43.76 ± 0.66 ** | 0.23 ± 0 ** |
| 5 | -1 | -1 | 1 | -1 | TS |  | 5.06 ± 0.07 ** | 40 ± 0.79 ** | 0.2 ± 0.01 ** |
| 6 | 1 | -1 | 1 | -1 | ACL_TS |  | 5.57 ± 0.08 ** | 45.37 ± 1.5 ** | 0.25 ± 0.01 ** |
| 7 | -1 | 1 | 1 | -1 | TS_ACC |  | 4.95 ± 0.21 * | 41.16 ± 0.06 ** | 0.22 ± 0 ** |
| 8 | 1 | 1 | 1 | -1 | ACL_ACC_TS |  | 5.52 ± 0.11 ** | 51.85 ± 1.25 ** | 0.29 ± 0.02 ** |
| 9 | -1 | -1 | -1 | 1 | WT | 175 | 5.65 ± 0.05 | 49.32 ± 0.88 | 0.24 ± 0 |
| 10 | 1 | -1 | -1 | 1 | ACL |  | 5.64 ± 0.08 | 50.34 ± 0.67 | 0.25 ± 0.01 |
| 11 | -1 | 1 | -1 | 1 | ACC |  | 5.58 ± 0.08 | 49.51 ± 0.62 | 0.25 ± 0.01 |
| 12 | 1 | 1 | -1 | 1 | ACL_ACC |  | 6.41 ± 0.16 ** | 59.45 ± 0.36 ** | 0.26 ± 0.01 ** |
| 13 | -1 | -1 | 1 | 1 | TS |  | 6.08 ± 0.07 ** | 49.81 ± 1.32 | 0.25 ± 0 |
| 14 | 1 | -1 | 1 | 1 | ACL_TS |  | 6.86 ± 0.16 ** | 59.78 ± 0.65 ** | 0.29 ± 0.01 ** |
| 15 | -1 | 1 | 1 | 1 | TS_ACC |  | 5.68 ± 0.13 | 58.04 ± 0.52 ** | 0.24 ± 0.01 |
| 16 | 1 | 1 | 1 | 1 | ACL_ACC_TS |  | 6.69 ± 0.15 ** | 54.66 ± 1.35 ** | 0.29 ± 0.01 ** |
| Differences with control (WT grown at the same C/N ratio medium) were evaluated using a t-test *: $p \leq 0.05$, **: $p \leq 0.01$. | | | | | | | | | |

**Figure S2.**
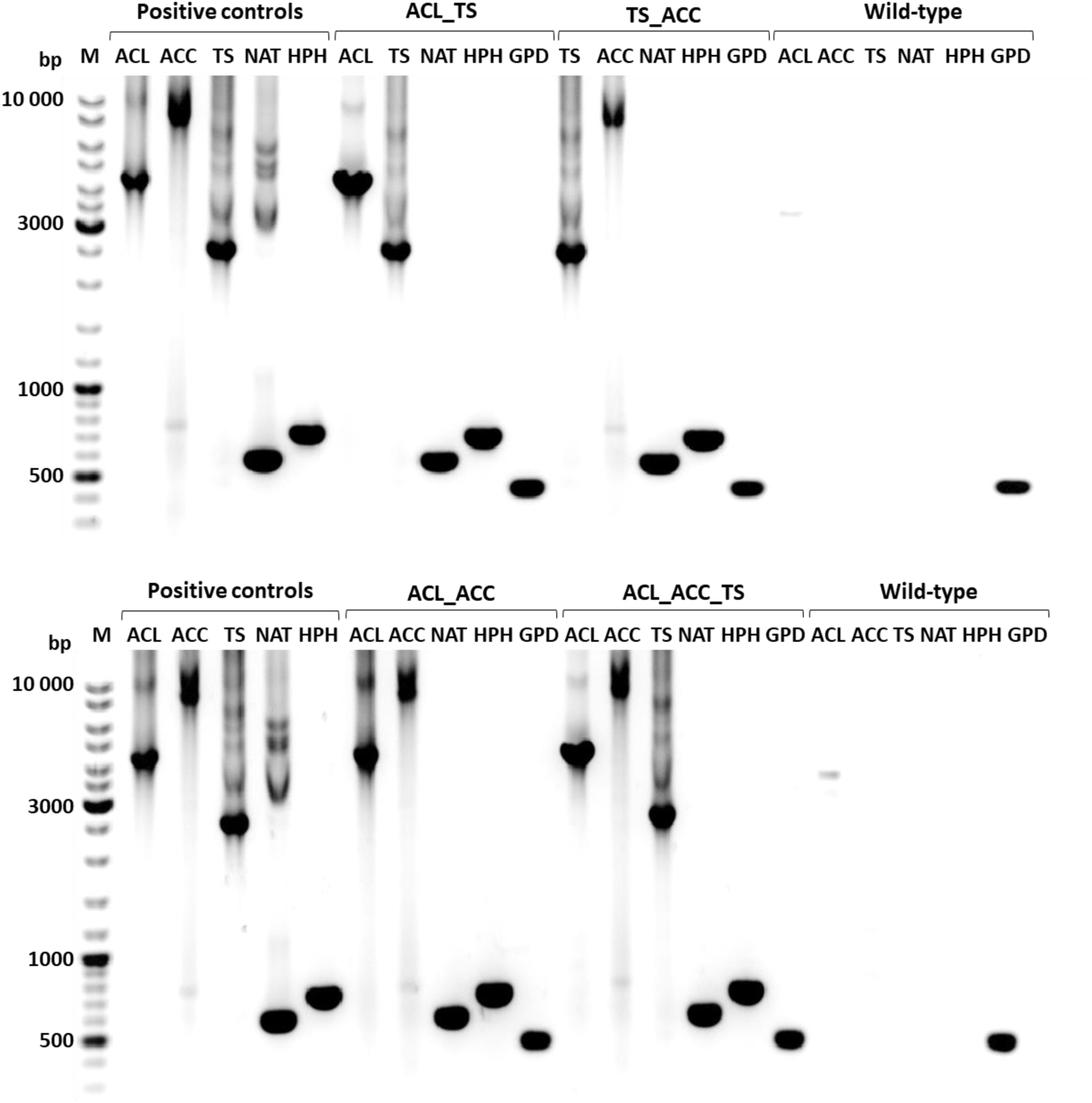
Colony PCR products were run on 1 % agarose gel. Positive controls were prepared by using corresponding primers and plasmids. The quality of gDNA in PCR reaction was assessed by using primers specific for glyceraldehyde-3-phosphate dehydrogenase gene (GPD). NAT: Nourseothricin acetyltransferase, HPT: Hygromycin B phosphotransferase, ACL: ATP-citrate lyase, ACC: acetyl-CoA carboxylase, TS: threonine synthase.

**Table S4.** Fatty acid profile of transformants grown at C/N30 and 175 minimal medium at 96h. The fatty acid profile of palm oil was added for direct comparison. MUFAs: Monounsaturated fatty acids, PUFAs: Polyunsaturated fatty acids. Differences with control (WT grown at the same C/N ratio medium) were evaluated using a t-test *: p ≤ 0.05, **: p ≤ 0.01.

| Fatty Acid Profile (%) |  |  |  |  |  |  |  |  |  |  |  |  |  |  |  |  |
| --- | --- | --- | --- | --- | --- | --- | --- | --- | --- | --- | --- | --- | --- | --- | --- | --- |
| C/N<br>(g/g) | Strains | C14:0 | C16:0 | C16:1 | C18:0 | C18:1 | C18:2 | C18:3 | C20:0 | C20:1 | C22:0 | C24:0 | Saturated<br>FAs | MUFAs | PUFAs | Very<br>Long<br>Chain<br>FAs |
| 30 | WT | 0.35 ±<br>0.02 | 22.28 ±<br>0.07 | 1.57 ±<br>0.10 | 12.90 ±<br>0.33 | 49.50 ±<br>0.16 | 10.45<br>± 0.76 | 0.29 ±<br>0.00 | 0.51 ±<br>0.08 | - | 0.31 ±<br>0.01 | 1.71 ±<br>0.09 | 38.21 ±<br>0.51 | 51.08 ±<br>0.24 | 10.71 ±<br>0.74 | 2.07 ±<br>0.14 |
|  | ACL | 0.32 ±<br>0.03 | 21.62 ±<br>0.78 | 1.56 ±<br>0.08 | 13.66 ±<br>0.36 | 50.61 ±<br>0.30 | 9.22 ±<br>0.16 | 0.28 ±<br>0.0 | 0.88 ±<br>0.14 | - | 0.44 ±<br>0.03 | 1.43 ±<br>0.02 | 38.36 ±<br>0.35 | 52.16 ±<br>0.22 ** | 9.48 ±<br>0.17 | 1.87 ±<br>0.02 |
|  | ACC | 0.36 ±<br>0.01 | 22.07 ±<br>0.76 | 1.63 ±<br>0.06 | 12.68 ±<br>0.37 | 50.97 ±<br>0.21 | 9.34 ±<br>0.58 | 0.27 ±<br>0.02 | 0.95 ±<br>0.02 | - | 0.42 ±<br>0.01 | 1.39 ±<br>0.15 | 37.79 ±<br>0.70 | 52.61 ±<br>0.15 ** | 9.61 ±<br>0.60 | 1.81 ±<br>0.15 |
|  | TS | 0.41 ±<br>0.02 | 20.95 ±<br>0.08 | 1.18 ±<br>0.05 | 17.76 ±<br>0.10 | 46.99 ±<br>0.04 | 8.32 ±<br>0.10 | 0.57 ±<br>0.03 | 1.06 ±<br>0.10 | - | 0.53 ±<br>0.01 | 2.22 ±<br>0.15 | 42.93 ±<br>0.07 ** | 48.18 ±<br>0.07 ** | 8.89 ±<br>0.11 * | 2.75 ±<br>0.16 * |
|  | ACL_TS | 0.41 ±<br>0.02 | 21.26 ±<br>1.05 | 0.68 ±<br>0.06 | 17.98 ±<br>0.50 | 46.18 ±<br>0.38 | 8.30 ±<br>0.48 | 0.95 ±<br>0.03 | 0.49 ±<br>0.02 | 1.98 ±<br>0.08 | 0.42 ±<br>0.01 | 1.82 ±<br>0.17 | 42.38 ±<br>0.70 ** | 48.84 ±<br>0.26 ** | 9.25 ±<br>0.51 | 2.25 ±<br>0.18 |
|  | ACL_ACC | 0.36 ±<br>0.04 | 18.89 ±<br>0.83 | 1.08 ±<br>0.08 | 21.01 ±<br>0.56 | 46.44 ±<br>0.30 | 8.05 ±<br>0.10 | 0.48 ±<br>0.03 | 0.62 ±<br>0.03 | 0.83 ±<br>0.02 | 0.53 ±<br>0.01 | 1.96 ±<br>0.15 | 43.36 ±<br>0.20 ** | 48.34 ±<br>0.24 ** | 8.52 ±<br>0.12 * | 2.49 ±<br>0.15 * |
|  | TS_ACC | 0.47 ±<br>0.01 | 21.90 ±<br>0.35 | 1.14 ±<br>0.05 | 18.31 ±<br>0.53 | 45.54 ±<br>0.37 | 8.11 ±<br>0.03 | 0.96 ±<br>0.05 | 0.56 ±<br>0.02 | 0.79 ±<br>0.06 | 0.50 ±<br>0.01 | 2.25 ±<br>0.15 | 43.97 ±<br>0.34 ** | 47.47 ±<br>0.39 ** | 9.07 ±<br>0.04 * | 2.75 ±<br>0.14 ** |
|  | ACL_ACC_TS | 0.36 ±<br>0.01 | 20.63 ±<br>0.66 | 0.99 ±<br>0.33 | 19.17 ±<br>1.35 | 45.51 ±<br>0.60 | 8.27 ±<br>0.44 | 0.48 ±<br>0.04 | 0.77 ±<br>0.04 | 1.29 ±<br>0.05 | 0.52 ±<br>0.03 | 2.23 ±<br>0.16 | 43.68 ±<br>1.26 ** | 47.79 ±<br>0.77 ** | 8.75 ±<br>0.45 * | 2.75 ±<br>0.18 ** |
| 175 | WT | 0.32 ±<br>0.01 | 21.53 ±<br>0.62 | 1.4 ±<br>0.12 | 13.63 ±<br>0.69 | 51.39 ±<br>0.24 | 8.29 ±<br>0.03 | 0.28 ±<br>0.01 | 1.15 ±<br>0.06 | - | 0.47 ±<br>0.02 | 1.42 ±<br>0.04 | 38.52 ±<br>0.31 | 52.79 ±<br>0.23 | 8.69 ±<br>0.18 | 1.89 ±<br>0.04 |
|  | ACL | 0.29 ±<br>0.01 | 21.02 ±<br>0.56 | 1.16 ±<br>0.11 | 14.70 ±<br>0.49 | 51.77 ±<br>0.53 | 7.75 ±<br>0.11 | 0.27 ±<br>0.01 | 1.17 ±<br>0.03 | - | 0.48 ±<br>0.02 | 1.38 ±<br>0.04 | 39.05 ±<br>0.56 | 52.93 ±<br>0.52 | 8.02 ±<br>0.11 * | 1.86 ±<br>0.04 |
|  | ACC | 0.31 ±<br>0.00 | 21.62 ±<br>0.47 | 1.34 ±<br>0.02 | 13.89 ±<br>0.34 | 51.84 ±<br>0.20 | 7.76 ±<br>0.08 | 0.28 ±<br>0.00 | 1.21 ±<br>0.05 | - | 0.48 ±<br>0.02 | 1.28 ±<br>0.06 | 38.79 ±<br>0.30 | 53.18 ±<br>0.23 | 8.03 ±<br>0.08 * | 1.76 ±<br>0.05 |
|  | TS | 0.28 ±<br>0.00 | 18.12 ±<br>0.62 | 0.75 ±<br>0.02 | 20.61 ±<br>0.74 | 49.70 ±<br>0.24 | 6.72 ±<br>0.15 | 0.48 ±<br>0.03 | 1.13 ±<br>0.04 | 0.11 ±<br>0.01 | 0.61 ±<br>0.03 | 1.47 ±<br>0.09 | 42.24 ±<br>0.34 ** | 50.56 ±<br>0.26 ** | 7.21 ±<br>0.12 ** | 2.09 ±<br>0.10 |
|  | **ACL_TS** | 0.24 ±<br>0.012 | 18.86 ±<br>0.88 | 0.66 ±<br>0.03 | 18.65 ±<br>0.51 | 48.77 ±<br>0.24 | 7.40 ±<br>0.19 | 0.81 ±<br>0.02 | 0.97 ±<br>0.02 | 1.21 ±<br>0.05 | 0.52 ±<br>0.02 | 1.90 ±<br>0.12 | 41.14 ±<br>0.39 ** | 50.64 ±<br>0.26 ** | 8.22 ±<br>0.20 | 2.42 ±<br>0.13 ** |
|  | **ACL_ACC** | 0.24 ±<br>0.00 | 17.12 ±<br>0.62 | 0.88 ±<br>0.03 | 20.10 ±<br>0.67 | 49.72 ±<br>0.25 | 7.44 ±<br>0.13 | - | 1.14 ±<br>0.02 | 0.97 ±<br>0.04 | 0.58 ±<br>0.02 | 1.81 ±<br>0.09 | 40.99 ±<br>0.26 ** | 51.57 ±<br>0.20 ** | 7.44 ±<br>0.13 ** | 2.39 ±<br>0.07 ** |
|  | **TS_ACC** | 0.24 ±<br>0.01 | 18.94 ±<br>1.82 | 0.96 ±<br>0.08 | 18.59 ±<br>1.17 | 48.28 ±<br>0.87 | 7.59 ±<br>0.46 | 0.65 ±<br>0.03 | 0.93 ±<br>0.06 | 1.39 ±<br>0.07 | 0.71 ±<br>0.01 | 1.84 ±<br>0.08 | 41.24 ±<br>0.54 ** | 50.63 ±<br>0.86 * | 8.23 ±<br>0.43 | 2.54 ±<br>0.09 ** |
|  | **ACL_ACC_TS** | 0.27 ±<br>0.01 | 20.59 ±<br>0.07 | 1.26 ±<br>0.14 | 15.04 ±<br>0.42 | 50.13 ±<br>0.39 | 8.35 ±<br>0.38 | 0.54 ±<br>0.02 | 0.76 ±<br>0.04 | 1.19 ±<br>0.04 | 0.47 ±<br>0.02 | 1.38 ±<br>0.07 | 38.52 ±<br>0.35 | 52.59 ±<br>0.30 | 8.89 ±<br>0.38 | 1.84 ±<br>0.05 |
|  | **Palm (range)**<br>**[13]** | 0.3 –<br>1.1 | 32.6–<br>46.0 | 0.0 –<br>0.5 | 1.7–7.6 | 35.1 –<br>53.4 | 7.0 –<br>15.0 | 0.2 –<br>0.6 | 0.1–0.5 | 0.1 –<br>0.2 | - | – | 36.7 –<br>51.5 | 35.3 –<br>53.8 | 7.3 –<br>15.5 | - |
| Differences with control (WT grown at the same C/N ratio medium) were evaluated using a t-test *: $p \leq 0.05$, **: $p \leq 0.01$. | | | | | | | | | | | | | | | | |

**Table S5.** Model coefficients, and statistics of linear regression model.

| Dependent variable: Total lipid (g/L) |  |  |  |  |
| --- | --- | --- | --- | --- |
| Source | Estimate | Std. Error | t value | p-value |
| Intercept | 2.68 | 0.0109 | 245.0180 | < e-16 |
| ACL | 0.301 | 0.0109 | 27.5490 | < e-16 |
| ACC | 0.113 | 0.0109 | 10.3320 | < e-11 |
| TS | 0.258 | 0.0109 | 23.6010 | < e-16 |
| CN | 0.602 | 0.0109 | 55.0500 | < e-16 |
| ACL:ACC | 0.047 | 0.0109 | 4.2940 | 0.00015 |
| ACL:TS | 0.045 | 0.0109 | 4.1140 | 0.00025 |
| ACC:TS | -0.094 | 0.0109 | -8.6340 | < e-10 |
| ACL:CN | 0.012 | 0.0109 | 1.0840 | 0.286 |
| ACC:CN | -0.014 | 0.0109 | -1.2920 | 0.320 |
| TS:CN | -0.021 | 0.0109 | -1.9210 | 0.06369 |
| ACL:ACC:TS | -0.094 | 0.0109 | -8.5880 | < e-10 |
| ACL:ACC:CN | -0.001 | 0.0109 | -0.7880 | 0.4422 |
| ACL:TS:CN | -0.002 | 0.0109 | -0.1730 | 0.86344 |
| ACC:TS:CN | -0.049 | 0.0109 | -4.5030 | < e-5 |
| ACL:ACC:TS:CN | -0.122 | 0.0109 | -11.1570 | < e-12 |
| R<sup>2</sup>: 0.9934, Adj R<sup>2</sup>:0.9903, p-value: < 2.2e-16 |  |  |  |  |

**Table S6.** Fatty acid profile of wild-type in the bioreactor experiment.

| Time<br>(h) | Fatty acid profile (%) |  |  |  |  |  |  |  |  |  |  |  |  |  |  |  |
| --- | --- | --- | --- | --- | --- | --- | --- | --- | --- | --- | --- | --- | --- | --- | --- | --- |
|  | C14:0 | C16:0 | C16:1 | C18:0 | C18:1 | C18:2 | C18:3 | C20:0 | C20:1 | C22:0 | C22:1 | C24:0 | Saturated<br>FA | MUFAs | PUFAs | VLC-FA |
| 17 | 0.85 ±<br>0.01 | 18.35<br>± 0.25 | 1.14 ±<br>0.02 | 6.50 ±<br>0.01 | 31.87<br>± 0.43 | 35.83<br>± 0.43 | 1.51 ±<br>0.39 | - | - | - | - | 2.93 ±<br>0.00 | 29.65 ±<br>0.41 | 33.02 ±<br>0.42 | 37.33 ±<br>0.00 | 2.93 ±<br>0.00 |
| 24 | 0.82 ±<br>0.01 | 19.44<br>± 0.12 | 1.25 ±<br>0.01 | 7.33 ±<br>0.00 | 32.38<br>± 0.39 | 35.16<br>± 0.39 | 1.08 ±<br>0.69 | - | - | - | - | 0.45 ±<br>0.03 | 30.12 ±<br>0.09 | 33.63 ±<br>0.64 | 36.25 ±<br>0.32 | 0.45 ±<br>0.19 |
| 48 | 0.76 ±<br>0.05 | 36.89<br>± 4.28 | 3.46 ±<br>0.53 | 7.68 ±<br>0.00 | 38.01<br>± 0.88 | 11.76<br>± 0.88 | 0.26 ±<br>3.27 | - | 0.30 ±<br>0.02 | - | 0.46 ±<br>0.00 | 0.49 ±<br>0.12 | 46.04 ±<br>0.36 | 42.23 ±<br>0.45 | 12.02 ±<br>0.45 | 0.95 ±<br>0.12 |
| 64 | 0.65 ±<br>0.03 | 32.76<br>± 0.98 | 2.97 ±<br>0.03 | 9.37 ±<br>0.00 | 42.10<br>± 0.57 | 10.14<br>± 0.57 | 0.22 ±<br>0.01 | 0.37 ±<br>0.04 | 0.85 ±<br>0.09 | - | 0.36 ±<br>0.00 | 0.70 ±<br>0.10 | 44.14 ±<br>0.71 | 46.28 ±<br>0.69 | 10.43 ±<br>0.35 | 1.06 ±<br>0.14 |
| 72 | 0.58 ±<br>0.05 | 30.60<br>± 1.46 | 2.73 ±<br>0.01 | 10.25<br>± 0.00 | 43.58<br>± 0.82 | 9.78 ±<br>0.82 | 0.30 ±<br>0.02 | 0.49 ±<br>0.00 | 0.30 ±<br>0.12 | 0.28 ±<br>0.03 | 0.50 ±<br>0.00 | 0.68 ±<br>0.01 | 42.96 ±<br>1.63 | 47.12 ±<br>0.93 | 10.22 ±<br>0.99 | 1.46 ±<br>0.02 |
| 96 | 0.46 ±<br>0.07 | 25.76<br>± 2.23 | 2.04 ±<br>0.06 | 14.05<br>± 0.00 | 45.87<br>± 0.91 | 8.56 ±<br>0.91 | 0.28 ±<br>0.04 | 0.90 ±<br>0.04 | 0.85 ±<br>0.08 | 0.53 ±<br>0.06 | - | 0.76 ±<br>0.01 | 42.54 ±<br>2.28 | 48.76 ±<br>0.93 | 8.98 ±<br>1.41 | 1.29 ±<br>0.05 |
| 120 | 0.43 ±<br>0.10 | 25.22<br>± 2.83 | 1.88 ±<br>0.19 | 13.52<br>± 0.00 | 47.20<br>± 1.44 | 8.70 ±<br>1.44 | 0.33 ±<br>0.01 | 0.94 ±<br>0.04 | 0.74 ±<br>0.11 | 0.52 ±<br>0.06 | 0.55 ±<br>0.00 | 0.50 ±<br>0.07 | 41.24 ±<br>2.79 | 50.36 ±<br>1.37 | 9.14 ±<br>1.39 | 1.56 ±<br>0.13 |
| 144 | 0.42 ±<br>0.07 | 25.24<br>± 1.83 | 1.85 ±<br>0.09 | 12.82<br>± 0.00 | 47.80<br>± 0.71 | 8.74 ±<br>0.70 | 0.35 ±<br>0.04 | 0.88 ±<br>0.04 | 0.53 ±<br>0.06 | 0.53 ±<br>0.06 | 0.48 ±<br>0.00 | 0.72 ±<br>0.01 | 40.67 ±<br>1.79 | 50.66 ±<br>0.67 | 9.19 ±<br>1.14 | 1.73 ±<br>0.08 |
| 168 | 0.40 ±<br>0.05 | 23.74<br>± 1.42 | 1.56 ±<br>0.06 | 13.85<br>± 0.00 | 48.58<br>± 0.32 | 8.45 ±<br>0.32 | 0.34 ±<br>0.05 | 0.95 ±<br>0.02 | 0.80 ±<br>0.08 | 0.54 ±<br>0.08 | 0.51 ±<br>0.00 | 0.88 ±<br>0.15 | 40.47 ±<br>1.11 | 51.45 ±<br>0.35 | 8.88 ±<br>0.40 | 1.93 ±<br>0.24 |
| 192 | 0.40 ±<br>0.04 | 23.49<br>± 1.01 | 1.53 ±<br>0.04 | 14.81<br>± 0.00 | 49.90<br>± 0.11 | 6.57 ±<br>0.11 | 0.14 ±<br>0.03 | 1.05 ±<br>0.05 | 1.51 ±<br>0.07 | 0.56 ±<br>0.08 | 0.86 ±<br>0.00 | 0.62 ±<br>0.10 | 41.01 ±<br>0.78 | 53.80 ±<br>0.09 | 6.70 ±<br>0.71 | 2.04 ±<br>0.19 |

**Table S7.** Fatty acid profile of ACL_ACC_TS in the bioreactor experiment.

| Time<br>(h) | Fatty acid profile (%) |  |  |  |  |  |  |  |  |  |  |  |  |  |  |  |
| --- | --- | --- | --- | --- | --- | --- | --- | --- | --- | --- | --- | --- | --- | --- | --- | --- |
|  | C14:0 | C16:0 | C16:1 | C18:0 | C18:1 | C18:2 | C18:3 | C20:0 | C20:1 | C22:0 | C22:1 | C24:0 | Saturated<br>FA | MUFAs | PUFAs | VLC-FA |
| 17 | 0.67 ±<br>0.04 | 16.81<br>± 0.41 | 0.93 ±<br>0.03 | 7.73 ±<br>0.10 | 31.09<br>± 0.17 | 38.41<br>± 0.52 | 2.83 ±<br>0.12 | - | - | - | - | 0.88 ±<br>0.03 | 26.75 ±<br>0.45 | 32.01 ±<br>0.20 | 41.23 ±<br>0.64 | 0.88 ±<br>0.03 |
| 24 | 0.57 ±<br>0.01 | 15.89<br>± 0.70 | 0.78 ±<br>0.00 | 12.59<br>± 0.21 | 35.46<br>± 0.09 | 30.03<br>± 0.01 | 2.37 ±<br>0.03 | - | - | - | - | 0.17 ±<br>0.36 | 31.07 ±<br>0.07 | 36.24 ±<br>0.08 | 32.69 ±<br>0.02 | 0.17 ±<br>0.56 |
| 48 | 0.44 ±<br>0.03 | 27.38<br>± 1.43 | 1.66 ±<br>0.16 | 15.04<br>± 1.24 | 42.27<br>± 0.66 | 10.08<br>± 0.38 | 0.92 ±<br>0.01 | 0.39 ±<br>0.04 | 0.64 ±<br>0.02 | 0.34 ±<br>0.03 | 0.40 ±<br>0.01 | 0.77 ±<br>0.00 | 44.57 ±<br>0.11 | 44.97 ±<br>0.45 | 11.10 ±<br>0.37 | 1.51 ±<br>0.02 |
| 64 | 0.48 ±<br>0.00 | 27.97<br>± 0.44 | 1.38 ±<br>0.13 | 14.86<br>± 0.33 | 43.34<br>± 0.17 | 8.97 ±<br>0.34 | 0.88 ±<br>0.12 | 0.39 ±<br>0.01 | 0.40 ±<br>0.02 | 0.34 ±<br>0.02 | 0.40 ±<br>0.02 | 0.68 ±<br>0.04 | 44.94 ±<br>0.15 | 45.52 ±<br>0.30 | 9.94 ±<br>0.48 | 1.42 ±<br>0.04 |
| 72 | 0.50 ±<br>0.01 | 29.02<br>± 0.38 | 1.44 ±<br>0.15 | 13.88<br>± 0.39 | 43.45<br>± 0.34 | 8.86 ±<br>0.32 | 0.81 ±<br>0.02 | 0.37 ±<br>0.04 | 0.62 ±<br>0.08 | 0.30 ±<br>0.00 | 0.44 ±<br>0.00 | 0.64 ±<br>0.04 | 44.91 ±<br>0.11 | 45.95 ±<br>0.28 | 9.75 ±<br>0.30 | 1.39 ±<br>0.05 |
| 96 | 0.51 ±<br>0.02 | 29.48<br>± 1.19 | 1.26 ±<br>0.12 | 13.54<br>± 1.03 | 44.28<br>± 0.19 | 8.24 ±<br>0.12 | 0.74 ±<br>0.09 | 0.36 ±<br>0.05 | 0.78 ±<br>0.14 | 0.31 ±<br>0.03 | 0.47 ±<br>0.01 | 0.54 ±<br>0.03 | 44.92 ±<br>0.04 | 46.80 ±<br>0.22 | 9.06 ±<br>0.03 | 1.32 ±<br>0.07 |
| 120 | 0.52 ±<br>0.02 | 29.18<br>± 1.26 | 1.16 ±<br>0.10 | 13.79<br>± 0.77 | 44.68<br>± 0.13 | 7.88 ±<br>0.24 | 0.76 ±<br>0.13 | 0.37 ±<br>0.05 | 0.78 ±<br>0.02 | 0.32 ±<br>0.04 | 0.50 ±<br>0.04 | 0.64 ±<br>0.00 | 44.94 ±<br>0.39 | 47.12 ±<br>0.00 | 8.71 ±<br>0.41 | 1.46 ±<br>0.00 |
| 144 | 0.51 ±<br>0.00= | 28.81<br>± 0.53 | 1.21 ±<br>0.02 | 13.21<br>± 0.03 | 45.61<br>± 0.05 | 7.96 ±<br>0.37 | 0.70 ±<br>0.06 | 0.39 ±<br>0.03 | 0.33 ±<br>0.04 | 0.32 ±<br>0.03 | 0.47 ±<br>0.03 | 0.67 ±<br>0.05 | 43.96 ±<br>0.50 | 47.62 ±<br>0.06 | 8.75 ±<br>0.40 | 1.46 ±<br>0.11 |
| 168 | 0.49 ±<br>0.00 | 27.87<br>± 0.05 | 1.16 ±<br>0.07 | 14.05<br>± 0.26 | 46.07<br>± 0.38 | 7.54 ±<br>0.37 | 0.58 ±<br>0.06 | 0.43 ±<br>0.00 | 0.75 ±<br>0.04 | 0.34 ±<br>0.04 | 0.54 ±<br>0.01 | 0.77 ±<br>0.02 | 44.08 ±<br>0.16 | 48.52 ±<br>0.58 | 8.15 ±<br>0.40 | 1.65 ±<br>0.24 |
| 192 | 0.49 ±<br>0.02 | 26.86<br>± 0.27 | 1.04 ±<br>0.07 | 15.09<br>± 0.65 | 47.21<br>± 0.39 | 6.55 ±<br>1.05 | 0.45 ±<br>0.15 | 0.45 ±<br>0.03 | 1.23 ±<br>0.28 | 0.39 ±<br>0.01 | 0.81 ±<br>0.10 | 0.53 ±<br>0.06 | 43.87 ±<br>0.78 | 50.29 ±<br>0.71 | 7.07 ±<br>1.22 | 1.73 ±<br>0.03 |

